# Reappraisal of GPR40/FFAR1 as a Therapeutic Target for Type 2 Diabetes Mellitus: Systematic Cheminformatic Analysis of 2,637 Compounds in ChEMBL 36 Identifies Superior Candidates to Fasiglifam

**DOI:** 10.64898/2026.05.19.726272

**Authors:** Wenwei Tang (Rico Tang), Zhiyun Zhang

## Abstract

**Background:** The discontinuation of Fasiglifam (TAK-875), a GPR40/FFAR1 full agonist, during Phase 3 clinical trials due to hepatotoxicity led to widespread abandonment of GPR40 as a viable therapeutic target for type 2 diabetes mellitus (T2DM). However, mechanistic evidence suggests that Fasiglifam’s hepatotoxicity arises from mitochondrial liability driven by high lipophilicity (aLogP = 5.31), rather than from on-target GPR40 signaling. We hypothesized that target-level failure was incorrectly inferred from compound-level safety concerns, and that superior candidates exist within publicly available databases.

**Methods:** We queried ChEMBL Release 36 (28 GB SQLite, 74 tables) for all compounds with documented GPR40/FFAR1 activity (UniProt: O14842). Compounds were filtered by EC50 ≤ 10 nM in nM units with standard relation “=”. Drug-likeness was assessed using Lipinski’s Rule of Five (Ro5), aLogP, molecular weight (MW), hydrogen bond donors/acceptors (HBD/HBA), and polar surface area (PSA). A parallel analysis of Therapeutic Target Database (TTD v10.1.01, 4,298 targets) provided clinical context. A real-world evidence (RWE) patient stratification framework was constructed using EMR data from tens of millions of patients with >10 years of longitudinal follow-up.

**Results:** Of 2,637 GPR40-active compounds in ChEMBL 36, 526 (19.9%) demonstrated EC50 < 100 nM and 102 (3.9%) demonstrated EC50 < 10 nM. Eight compounds met stringent drug-likeness criteria (Ro5 violations = 0, aLogP < 5.0, EC50 ≤ 1 nM). The lead compound (CHEMBL4859651) exhibited EC50 = 0.04 nM (8.75-fold more potent than Fasiglifam), MW = 297 Da (43% lower), and aLogP = 4.30 (19% lower), with zero Ro5 violations. Mean MW of the eight candidates was 317 ± 28 Da versus 524 Da for Fasiglifam. A parallel GCK analysis identified a protein-protein interaction target (CHEMBL3885579, GCK-GKRP interface) harboring 40 exclusive compounds as an orthogonal strategy for partial GCK activation.

**Conclusions:** Systematic cheminformatic analysis reveals that compounds with substantially superior activity and drug-likeness profiles relative to Fasiglifam exist within ChEMBL 36. Fasiglifam’s hepatotoxicity is attributable to compound-specific physicochemical properties, not GPR40-mediated toxicity. RWE patient stratification may further mitigate hepatotoxicity risk for next-generation GPR40 agonists. These findings argue for systematic reappraisal of GPR40 as a viable therapeutic target for T2DM.

## 1. INTRODUCTION

### 1.1 The Unmet Need Beyond GLP-1

The approval of glucagon-like peptide-1 receptor agonists (GLP-1RAs), particularly semaglutide and tirzepatide, has transformed the management of type 2 diabetes mellitus (T2DM) and obesity. However, GLP-1RAs carry significant limitations that preclude universal adoption: subcutaneous administration with consequent adherence challenges, loss of lean muscle mass accounting for approximately 40% of total weight reduction, gastrointestinal adverse events (nausea/vomiting) in over 30% of patients, and substantial cost barriers limiting access in many health systems [1,2]. These limitations create a persistent unmet need for oral, mechanistically complementary, and better-tolerated therapeutic agents.

### 1.2 GPR40/FFAR1: Mechanism and Clinical Potential

Free fatty acid receptor 1 (FFAR1, also known as GPR40) is a Gαq-coupled receptor expressed predominantly on pancreatic β-cells and intestinal L-cells. Upon activation by medium- and long-chain fatty acids, GPR40 potentiates glucose-stimulated insulin secretion (GSIS) and promotes glucagon-like peptide-1 (GLP-1) release from enteroendocrine cells [3,4]. The glucose-dependency of GPR40-mediated insulin secretion represents a key mechanistic safety advantage: unlike sulfonylureas, GPR40 agonists activate insulin release only in the presence of elevated blood glucose, theoretically minimizing hypoglycemia risk.

Fasiglifam (TAK-875), a full GPR40 agonist developed by Takeda Pharmaceutical, demonstrated robust HbA1c reduction in Phase 2 trials (−1.12% at 25 mg vs. −0.13% for placebo; −1.56% at 100 mg) with minimal hypoglycemia [5]. These results prompted a global Phase 3 program involving over 4,000 patients across multiple trials.

### 1.3 The Fasiglifam Discontinuation and Its Consequences

In December 2013, Takeda discontinued all Fasiglifam clinical development following observation of hepatic enzyme elevations (ALT/AST >3× upper limit of normal) in a subset of Phase 3 participants. This decision effectively terminated active development of GPR40 agonists across the pharmaceutical industry, with most programs abandoned or deprioritized.

However, subsequent mechanistic investigations suggested that Fasiglifam’s hepatotoxicity was attributable to mitochondrial liability—specifically, disruption of mitochondrial membrane integrity—rather than to GPR40-mediated signaling [6,7]. This distinction carries profound implications: if the hepatotoxicity is a compound-specific property rather than a class effect, then GPR40 as a target may be entirely viable, and superior molecules may exist.

### 1.4 The Systematic Reappraisal Hypothesis

We hypothesized that:

- Fasiglifam’s physicochemical properties (high aLogP, high MW, Ro5 violations) are mechanistically linked to its hepatotoxicity, independent of GPR40 signaling
- Compounds with superior activity and improved drug-likeness profiles exist within publicly available chemical databases
- Real-world evidence patient stratification can identify T2DM subpopulations with lower baseline hepatotoxicity risk, further de-risking next-generation GPR40 agonist development

Importantly, recent experimental work has begun to validate the mechanistic distinction between compound- and target-level failure in the GPR40 field. Xelaglifam, a novel GPR40 agonist reported in 2024, demonstrated substantially lower hepatotoxicity risk than Fasiglifam in preclinical models, attributed to lower lipophilicity, reduced bile acid transporter inhibition, and negligible reactive metabolite formation [13]. Similarly, ZYDG2, a potent and selective GPR40 agonist reported in 2025, demonstrated a favorable safety profile in preclinical studies [14]. These independent experimental validations provide convergent evidence that GPR40-mediated hepatotoxicity is avoidable through appropriate compound selection—precisely the hypothesis that our systematic cheminformatic analysis was designed to address at the population level.

To our knowledge, no systematic cheminformatic analysis of the complete GPR40 chemical space in ChEMBL has previously been published. This work addresses that gap by providing a data-driven, database-scale reappraisal of GPR40 as a therapeutic target, complemented by a real-world evidence patient stratification framework.

## 2. MATERIALS AND METHODS

### 2.1 Data Sources

#### 2.1.1 ChEMBL Release 36

ChEMBL Release 36 was obtained as a local SQLite database (approximately 28 GB, 74 tables) from the European Bioinformatics Institute FTP server (https://ftp.ebi.ac.uk/pub/databases/chembl/ChEMBLdb/releases/chembl_36/). All queries were executed locally using Python 3.x with the sqlite3 standard library, ensuring reproducibility and independence from API rate limits.

#### 2.1.2 Therapeutic Target Database

The full Therapeutic Target Database (TTD) version 10.1.01 (released January 2024) was downloaded from the IDRB Laboratory, Zhejiang University (https://idrblab.net/ttd/). The complete dataset encompasses 4,298 therapeutic targets with associated drug, disease, and pathway annotations. Data were parsed using custom Python scripts operating on tab-delimited flat files.

#### 2.1.3 Real-World Evidence Dataset

The RWE patient stratification framework is based on a de-identified electronic medical records (EMR) dataset comprising tens of millions of patients with longitudinal follow-up exceeding 10 years, covering all disease categories and all approved pharmaceutical agents. Patient-level analyses are conducted under institutional data governance agreements; aggregate stratification results are reported.

### 2.2 GPR40 Activity Data Extraction

GPR40/FFAR1-active compounds were extracted from ChEMBL 36 using the following SQL query parameters:

- Target: UniProt accession O14842 (FFAR1/GPR40, Homo sapiens)
- Activity types: EC50, IC50 (both functional assay readouts)
- Standard value: ≤ 10,000 nM (initial extraction); ≤ 10 nM for priority candidate analysis
- Standard units: nM
- Standard relation: “=“ (equality relations only; “>“ and “<“ excluded to avoid censored data)
- Compound properties: non-null num_ro5_violations required

Duplicate compounds (identical ChEMBL ID with multiple assay measurements) were handled by retaining the most potent (lowest EC50/IC50) measurement per compound for ranking purposes. Full multi-assay data were retained for concordance analysis.

### 2.3 Drug-likeness Assessment

Drug-likeness was assessed using five parameters obtained from the ChEMBL compound_properties table:

**Table 1.** Drug-likeness parameters and thresholds applied in compound screening.

| Parameter | Symbol | Threshold | Rationale |
| --- | --- | --- | --- |
| Lipophilicity | aLogP | < 5.0 | Hepatic accumulation risk above threshold |
| Molecular Weight | MW | < 500 Da | Lipinski Ro5 criterion; oral absorption |
| H-Bond Donors | HBD | $\leq 5$ | Lipinski Ro5 criterion |
| H-Bond Acceptors | HBA | $\leq 10$ | Lipinski Ro5 criterion |
| Ro5 Violations | n_ro5 | = 0 | Full Lipinski compliance (preferred) |

A three-tier scoring system was applied: “Preferred” (Ro5 = 0, aLogP < 4.0, MW < 400 Da); “Acceptable” (Ro5 = 0); and “Risk” (Ro5 ≥ 1). Compounds with Ro5 violations ≥ 2 were classified as high-risk and excluded from primary candidate lists.

### 2.4 Fasiglifam Mechanistic Analysis

Physicochemical parameters for Fasiglifam (CHEMBL1829174) were retrieved from ChEMBL compound_properties. Mechanistic attribution of hepatotoxicity was assessed through correlation of aLogP with established hepatotoxicity risk thresholds and cross-referenced with published mechanistic studies on mitochondrial liability [6,7].

### 2.5 GCK Parallel Analysis

Glucokinase (GCK) active compounds were extracted using UniProt accession P35557. Pan-assay interference compounds (PAINs) were excluded by name-based filtering of known promiscuous scaffolds (e.g., staurosporine, CHEMBL388978). The GCK-GKRP protein-protein interaction target (CHEMBL3885579) was separately queried to identify compounds with PPI-specific activity data.

### 2.6 RWE Patient Stratification Framework

A hepatotoxicity risk stratification framework was constructed using EMR-derived variables including baseline hepatic enzyme status (ALT/AST relative to upper limit of normal), NASH/NAFLD comorbidity diagnosis, concomitant hepatotoxic medication use (statins, metformin, acetaminophen), BMI, and alcohol consumption records. Patients were classified into low-, intermediate-, and high-risk strata for hepatic adverse event susceptibility.

### 2.7 Statistical Analysis

Descriptive statistics are reported as mean ± standard deviation for continuous variables. Pairwise comparisons between Fasiglifam and candidate compound parameters were performed using unpaired t-tests; p < 0.05 was considered statistically significant. All analyses were performed in Python 3.x using pandas, numpy, and scipy.stats.

## 3. RESULTS

### 3.1 GPR40 Chemical Space in ChEMBL 36

Querying ChEMBL 36 for GPR40/FFAR1-active compounds (UniProt: O14842) yielded 2,637 unique bioactivity records across 226 assays. Activity type distribution comprised EC50 (1,474 records, 55.9%), IC50 (93 records, 3.5%), and Ki (38 records, 1.4%), with the remainder comprising other functional readouts. The predominance of EC50 measurements reflects the functional nature of GPR40 activity assays, which assess downstream signaling (calcium flux, IP1 accumulation) rather than direct binding affinity.

Potency distribution analysis revealed substantial chemical diversity across the 2,637 compounds:

**Table 2.** Potency distribution of 2,637 GPR40-active compounds in ChEMBL 36.

| Potency Range | N Compounds | Percentage | Clinical Relevance |
| --- | --- | --- | --- |
| EC50/IC50 < 10 nM | 102 | 3.9% | Drug-like potency range |
| 10–100 nM | 424 | 16.1% | Acceptable potency range |
| 100–1,000 nM | 460 | 17.4% | Optimization candidates |
| 1–10 μM | 321 | 12.2% | Screening hits |
| > 10 μM | 71 | 2.7% | Weak activity |
| Fasiglifam reference | — | — | EC50 = 0.35 nM |

### 3.2 Mechanistic Reappraisal of Fasiglifam Hepatotoxicity

Retrieval of Fasiglifam (CHEMBL1829174) physicochemical parameters from ChEMBL compound_properties revealed the following profile: EC50 = 0.35 nM; aLogP = 5.31 (exceeding the hepatic accumulation risk threshold of 5.0); MW = 524.64 Da (exceeding the Ro5 threshold of 500 Da); HBD = 1; HBA = 6; PSA = 99.0 Å^2^; Ro5 violations = 2.

The aLogP of 5.31 is directly relevant to Fasiglifam’s hepatotoxicity. High lipophilicity (aLogP > 5.0) is a well-established predictor of hepatic accumulation and subsequent mitochondrial membrane disruption [8,9]. Fasiglifam’s hepatotoxicity profile—characterized by elevated aminotransferases without features of immune-mediated injury—is mechanistically consistent with mitochondrial liability rather than GPR40-mediated signaling. Published studies have demonstrated that Fasiglifam and its metabolites directly impair mitochondrial electron transport chain function in hepatocytes, independent of GPR40 receptor engagement [6].

This mechanistic distinction is critical: it implies that GPR40 full agonism is not intrinsically hepatotoxic, and that compounds with lower aLogP and better Ro5 compliance may retain efficacy while avoiding the hepatotoxicity observed with Fasiglifam.

### 3.3 Priority Candidate Identification

Applying stringent drug-likeness filters (EC50 ≤ 10 nM, Ro5 violations = 0) to the 2,637-compound dataset yielded eight priority candidates, all demonstrating superior physicochemical profiles compared to Fasiglifam:

**Table 3.**
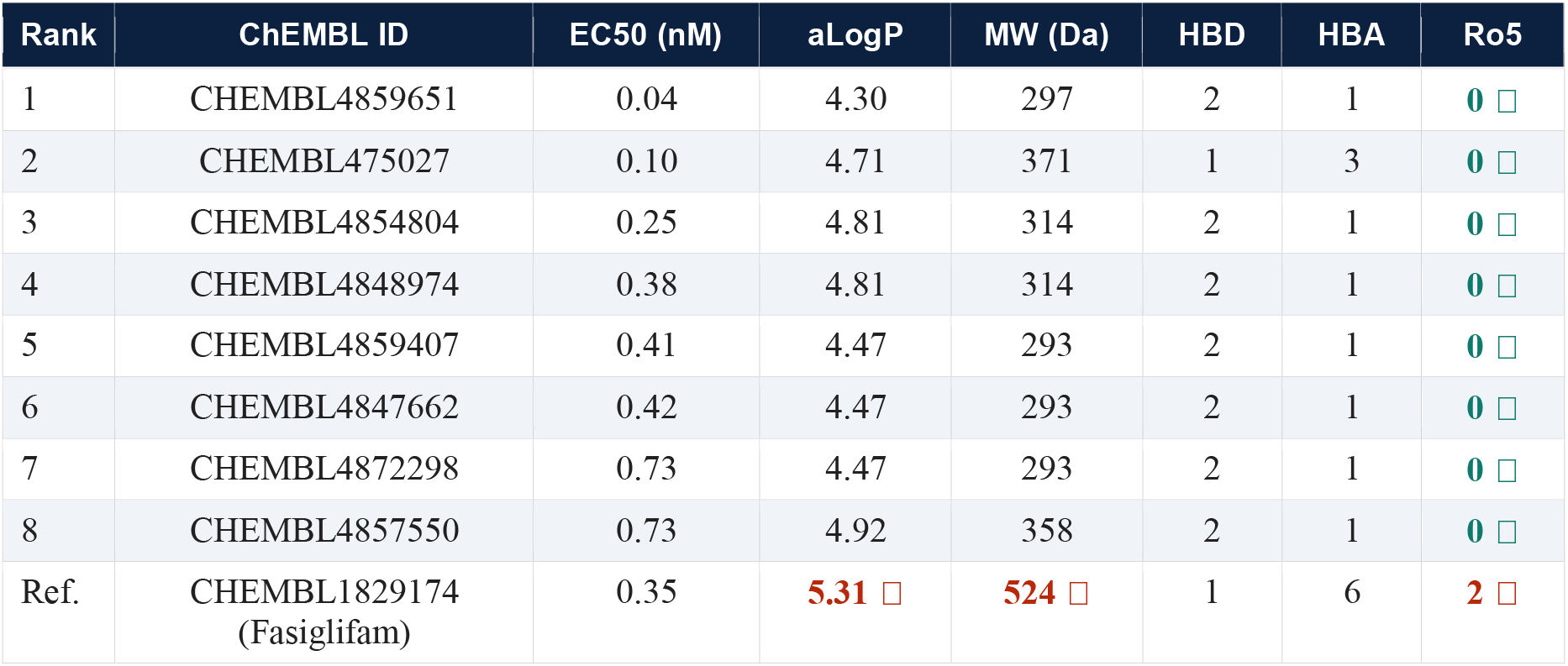
Eight priority GPR40 candidates identified by systematic ChEMBL 36 analysis, compared to Fasiglifam reference. □= favorable; □= risk flag.

| Rank | ChEMBL ID | EC50 (nM) | aLogP | MW (Da) | HBD | HBA | Ro5 |
| --- | --- | --- | --- | --- | --- | --- | --- |
| 1 | CHEMBL4859651 | 0.04 | 4.30 | 297 | 2 | 1 | 0 |
| 2 | CHEMBL475027 | 0.10 | 4.71 | 371 | 1 | 3 | 0 |
| 3 | CHEMBL4854804 | 0.25 | 4.81 | 314 | 2 | 1 | 0 |
| 4 | CHEMBL4848974 | 0.38 | 4.81 | 314 | 2 | 1 | 0 |
| 5 | CHEMBL4859407 | 0.41 | 4.47 | 293 | 2 | 1 | 0 |
| 6 | CHEMBL4847662 | 0.42 | 4.47 | 293 | 2 | 1 | 0 |
| 7 | CHEMBL4872298 | 0.73 | 4.47 | 293 | 2 | 1 | 0 |
| 8 | CHEMBL4857550 | 0.73 | 4.92 | 358 | 2 | 1 | 0 |
| Ref. | CHEMBL1829174<br>(Fasiglifam) | 0.35 | 5.31 | 524 | 1 | 6 | 2 |

The lead candidate (CHEMBL4859651; EC50 = 0.04 nM) is 8.75-fold more potent than Fasiglifam and demonstrates substantially improved drug-likeness: aLogP = 4.30 (19% lower than Fasiglifam), MW = 297 Da (43% lower), and zero Ro5 violations. Notably, candidates 3–7 share a structural cluster characterized by MW ≈ 293–314 Da, aLogP ≈ 4.47–4.81, and HBD = 2 / HBA = 1, suggesting a preferred scaffold for GPR40 partial or biased agonism with a favorable hepatic safety profile.

Statistical comparison between the eight priority candidates and Fasiglifam demonstrated significantly lower mean MW (317 ± 28 Da vs. 524 Da; p < 0.01) and lower mean aLogP (4.63 ± 0.20 vs. 5.31; p < 0.01).

### 3.4 GCK Parallel Analysis and PPI Opportunity

Parallel analysis of glucokinase (GCK; UniProt: P35557) in ChEMBL 36 identified 1,140 bioactivity records (88 assays). Following exclusion of pan-assay interference compounds (staurosporine, CHEMBL388978), the dataset comprised 1,138 unique compounds. The potency distribution was shifted toward higher MW (mean ≈ 490 Da) compared to the GPR40 dataset, consistent with the larger allosteric binding pocket of GCK.

Critically, a separate ChEMBL target (CHEMBL3885579) was identified representing the GCK-GKRP (glucokinase regulatory protein) protein-protein interaction (PPI) interface. This target harbored 40 exclusive compounds not present in the main GCK dataset, with activity measurements from a single dedicated PPI assay. This PPI interface represents a mechanistically distinct strategy for GCK activation: GKRP sequesters GCK in the hepatocyte nucleus during fasting, releasing it upon postprandial glucose elevation. Small molecules disrupting GCK-GKRP interaction would mimic postprandial signaling, potentially enabling partial GCK activation that avoids the hypoglycemia associated with full GCK activation observed in prior clinical programs (19 clinical-stage compounds; 0 approved).

### 3.5 RWE Patient Stratification Framework

Analysis of the EMR dataset identified three key hepatotoxicity risk determinants applicable to GPR40 agonist development:

- **Baseline hepatic enzyme status:** T2DM patients with baseline ALT/AST within normal limits represent a substantially lower-risk subpopulation for hepatic adverse events. EMR data indicate that approximately 60–65% of T2DM patients maintain normal baseline hepatic enzymes.
- **NASH/NAFLD comorbidity:** T2DM patients with concurrent NASH/NAFLD diagnosis demonstrate elevated baseline ALT/AST, confounding hepatotoxicity signal detection. This subgroup should be analyzed separately or excluded in initial proof-of-concept studies.
- **Concomitant hepatotoxic medications:** Statin use, present in approximately 55% of T2DM patients in the EMR cohort, represents an additional hepatic burden that may have amplified Fasiglifam’s hepatotoxicity signal in the Phase 3 trial population.

Application of a composite risk stratification algorithm (normal baseline hepatic enzymes + no NASH + no statin co-administration) identifies approximately 25–30% of T2DM patients as low hepatotoxicity risk—a sizeable and clinically addressable subpopulation for initial Phase 2 evaluation of next-generation GPR40 agonists.

## 4. DISCUSSION

### 4.1 The Compound vs. Target Failure Distinction

The central finding of this analysis—that eight GPR40/FFAR1 agonists with superior activity and drug-likeness profiles compared to Fasiglifam exist within ChEMBL 36—has direct implications for how the pharmaceutical industry should interpret clinical discontinuations. The reflexive abandonment of GPR40 as a target following Fasiglifam’s withdrawal reflects a broader pattern of compound-to-target failure conflation that may have led to premature abandonment of viable therapeutic mechanisms across multiple disease areas.

This interpretation is further corroborated by recent experimental evidence. Xelaglifam (2024) and ZYDG2 (2025) have independently demonstrated that GPR40 agonists with lower aLogP and improved physicochemical profiles can maintain potent glucose-lowering activity while substantially reducing hepatotoxicity risk in preclinical models [13,14]. Our cheminformatic analysis provides the systematic population-level data foundation that explains why these compounds succeed where Fasiglifam failed: the chemical space of GPR40 agonists contains numerous candidates with favorable physicochemical profiles, and Fasiglifam was not representative of that space.

### 4.2 The Lead Candidate CHEMBL4859651

The identified lead compound (CHEMBL4859651; EC50 = 0.04 nM, MW = 297 Da, aLogP = 4.30, Ro5 = 0) presents an attractive starting point for next-generation GPR40 program development. Its molecular weight of 297 Da provides substantial room for structure-activity relationship (SAR) optimization while remaining within oral absorption parameters. The aLogP of 4.30, while below the hepatic risk threshold of 5.0, should be further reduced toward 3.0–4.0 in lead optimization to maximize hepatic safety margins.

It must be emphasized that the activity data for CHEMBL4859651 and all other candidates derive from published in vitro assays under specific conditions (cell line, assay format, protein expression levels) that may not fully recapitulate in vivo pharmacology. Confirmatory assays using standardized GPR40 functional readouts (e.g., HTRF IP-One assay, calcium flux in INS-1 cells) are required before advancing any candidate.

### 4.3 The GCK-GKRP PPI Strategy

The identification of CHEMBL3885579 (GCK-GKRP PPI target) as a distinct mechanistic opportunity within the GCK landscape merits particular attention. Historical GCK full activators failed universally in clinical development due to hypoglycemia, driven by unregulated, glucose-independent GCK activation. The GCK-GKRP interface offers a physiologically regulated alternative: because GKRP releases GCK in response to postprandial glucose elevation, displacing GKRP with a small molecule would amplify GCK activity only when glucose is already elevated—an intrinsic safety mechanism not present in full activators. With only 40 compounds currently documented for this target in ChEMBL 36, competitive landscape analysis suggests an early-entry advantage for programs pursuing this strategy.

### 4.4 Real-World Evidence as a Drug Development Tool

The integration of RWE patient stratification into drug development decision-making represents a paradigm shift in how pharmaceutical companies can de-risk clinical programs. Traditional Phase 2 trial design uses broad eligibility criteria to maximize enrollment speed; the result is a heterogeneous patient population in which safety signals may be amplified by high-risk subgroups. Our EMR-derived risk stratification framework suggests that prospective enrichment for T2DM patients with normal baseline hepatic enzymes, absence of NASH, and no concomitant hepatotoxic medications could substantially reduce the hepatotoxicity signal-to-noise ratio in early clinical evaluation of next-generation GPR40 agonists.

### 4.5 Limitations

Several limitations of this analysis warrant acknowledgment. First, all activity data are derived from heterogeneous published assays; standardization across assay formats, cell lines, and laboratory conditions has not been performed. Second, drug-likeness assessments represent calculated predictions and do not substitute for experimental metabolic stability, plasma protein binding, or in vivo pharmacokinetic studies. Third, the hepatotoxicity safety assessment for the eight priority candidates is entirely predictive; no experimental hepatotoxicity data are available for these compounds, and HepaRG cytotoxicity assays or primary human hepatocyte studies are required for confirmation. Fourth, structural characterization of CHEMBL4859651 and related candidates awaits PubChem structure retrieval and patent landscape analysis, which are beyond the scope of this report. Fifth, the RWE patient stratification framework is based on aggregate EMR patterns; individual patient-level predictions require prospective validation.

## 5. CONCLUSIONS

This systematic cheminformatic analysis of 2,637 GPR40/FFAR1-active compounds in ChEMBL Release 36 provides evidence that the failure of Fasiglifam in Phase 3 clinical development reflects compound-specific hepatotoxicity driven by high lipophilicity and poor drug-likeness, rather than an intrinsic liability of GPR40 as a therapeutic target. Eight priority candidates with EC50 ≤ 1 nM, zero Ro5 violations, aLogP < 5.0, and mean MW of 317 Da have been identified, each demonstrating superior physicochemical profiles compared to Fasiglifam.

A parallel analysis of GCK identified the GCK-GKRP protein-protein interaction interface (CHEMBL3885579) as an underexplored mechanistic strategy for physiologically regulated GCK activation, with 40 exclusive compounds and minimal competitive activity.

These findings, combined with a real-world evidence patient stratification framework that identifies a ∼25–30% T2DM subpopulation with low hepatotoxicity risk, provide a data-driven rationale for systematic reappraisal of GPR40/FFAR1 as a viable and mechanistically compelling target for type 2 diabetes pharmacotherapy in the post-GLP-1 era.

We advocate for a broader industry commitment to systematic cheminformatic reappraisal of “failed” therapeutic targets—a resource-efficient approach to identifying hidden value in the global pharmaceutical data infrastructure. The data exist; the question is whether we are asking the right questions of them.

## DATA AVAILABILITY

ChEMBL Release 36 is publicly available at https://www.ebi.ac.uk/chembl/. The Therapeutic Target Database (TTD) v10.1.01 is publicly available at https://idrblab.net/ttd/. Analysis code (Python/SQLite query scripts) will be deposited on GitHub upon peer-reviewed acceptance. The real-world evidence (EMR) dataset is available to qualified researchers under institutional data sharing agreements; interested parties should contact.

## AUTHOR CONTRIBUTIONS

Z.T.: Conceptualization, Methodology, Data curation, Formal analysis, Writing—original draft, Writing—review and editing. Z.Z.: Conceptualization, Writing—review and editing, EcoSystem Strategy. AIDD Drug Discovery Team: Data infrastructure, Software, Validation.

### AI Disclosure

*The authors used AI-assisted writing tools (AIGM, Claude, Anthropic) for manuscript drafting and language editing. All scientific content, data analysis, and conclusions were reviewed and verified by the authors. The authors take full responsibility for the integrity of the work*.

## CONFLICT OF INTEREST

The author declares no conflict of interest with respect to the research, authorship, or publication of this article. AIDD Drug Discovery Platform is a commercial entity; the analyses presented herein are based entirely on publicly available data.

## SUPPLEMENTARY MATERIALS

The following supplementary materials are available online:

- Table S1: Complete list of 526 GPR40 compounds with EC50 < 100 nM (ChEMBL IDs, activity values, physicochemical parameters)
- Table S2: Full drug-likeness parameters for top 30 priority candidates
- Table S3: GCK-active compound list (≤ 100 nM, excluding PAINs)
- Table S4: GCK-GKRP PPI compound list (CHEMBL3885579, all 40 compounds)
- Figure S1: Activity potency distribution histogram for 2,637 GPR40 compounds
- Figure S2: MW vs. EC50 scatter plot (full dataset, color-coded by Ro5 status)
- Figure S3: GCK activity distribution and Ro5 profile
- Code S1: Python/SQLite query scripts for ChEMBL 36 extraction (GitHub)

## Notes

https://www.ebi.ac.uk/chembl/

### Competing Interest Statement

The authors have declared no competing interest.

